# Cap analysis of gene expression (CAGE) sequencing reveals alternative promoter usage in complex disease

**DOI:** 10.1101/2021.08.28.458014

**Authors:** Sonal Dahale, Jorge Ruiz-Orera, Jan Silhavy, Norbert Hubner, Sebastiaan van Heesch, Michal Pravenec, Santosh S Atanur

## Abstract

The role of alternative promoter usage in tissue specific gene expression has been well established, however, its role in complex diseases is poorly understood. We performed cap analysis of gene expression (CAGE) tag sequencing from the left ventricle (LV) of a rat model of hypertension, the spontaneously hypertensive rat (SHR), and a normotensive strain, the Brown Norway (BN) to understand role of alternative promoter usage in complex disease. We identified 26,560 CAGE-defined transcription start sites (TSS) in the rat LV, including 1,970 novel cardiac TSS resulting in new transcripts. We identified 27 genes with alternative promoter usage between SHR and BN which could lead to protein isoforms differing at the amino terminus between two strains. Additionally, we identified 475 promoter switching events where a shift in TSS usage was within 100bp between SHR and BN, altering length of the 5’ UTR. Genomic variants located in the shifting promoter regions showed significant allelic imbalance in F1 crosses, confirming promoter shift. We found that the insulin receptor gene (*Insr*) showed a switch in promoter usage between SHR and BN in heart and liver. The *Insr* promoter shift was significantly associated with insulin levels and blood pressure within a panel of BXH/HXB recombinant inbred (RI) rat strains. This suggests that the hyperinsulinemia due to insulin resistance might lead to hypertension in SHR. Our study provides a preliminary evidence of alternative promoter usage in complex diseases.

## Introduction

Mammalian gene transcription is tightly regulated. The core promoter, defined as a ~40bp upstream and the downstream of the transcription start site (TSS) is sufficient for initiation of the transcription^1^. The general transcription initiation factors assemble at the core promoter with RNA Polymerase II to form a multi-protein complex that facilitates accurate transcription^2^. In mammalian genomes, although the number of protein coding genes is limited, the transcript repertoire is much more diverse^3^. It has been estimated that more than 60% of the mammalian genes have multiple transcripts^4^. In part, the transcriptional diversity is achieved through use of alternate promoters and alternate splicing^3^. The majority of the mammalian protein coding genes are regulated by multiple promoters that initiate transcription for multiple gene isoforms^4,5^. Alternate splicing regulates gene isoform expression post transcriptionally. On the contrary, alternate promoters provide a way to regulate gene isoform expression pre transcriptionally^6,7^.

The transcriptional diversity due to alternate promoters could be assessed by precisely mapping 5’ ends of the transcripts. Cap Analysis of Gene Regulation (CAGE), which takes advantage of the 7-methylguanosine cap structure of the transcripts, allows precise genome wide mapping of TSS at single base pair resolution^5,8^. Using CAGE, the FANTOM consortium has mapped precise TSSs across multiple tissues and primary cells from mouse and humans^4^. Approximately, 80% of the CAGE defined TSS showed tissue specific expression, suggesting that the majority of mammalian genes use alternative promoters in tissue specific manner to regulate tissue specific gene expression^4^. Furthermore, it has been shown that the transcripts switch promoters between oocytes and the zygotic stage during embryonic development in Zebrafish^9^. Often this shift in TSS usage is within 100bp of the same promoter. Though alternative promoter usage between different tissues and cell types, and between various developmental stages is well established^4,9^, the role of alternate promoter usage in disease has not been studied extensively.

The spontaneously hypertensive rat (SHR) strain is one of the most widely used animal models to study hypertension, which also shows many metabolic phenotypes, including insulin resistance, dyslipidaemia and central obesity, collectively known as metabolic syndrome (MtS)^10^. By crossing SHR with the Brown Norway (BN) strain, the BXH/HXB panel of recombinant inbreed (RI) rat strains have been derived^11^. Large numbers of physiological quantitative trait loci (pQTLs), expression QTLs (eQTLs) and histone QTLs (hQTLs) have been mapped using RI strains^12–16^. Whole genome sequencing of both the parental strains SHR and BN have revealed more than 4 million genomic variants between the two strains^10,17,18^. Availability of a rich resource of phenotypic and genotypic data, and homozygosity throughout the genome provides a unique opportunity to perform genetic studies in SHR.

To understand the role of alternative promoter usage in complex disease, we performed cap analysis of gene expression (CAGE) tag sequencing from left ventricles of SHR and BN rat strains. Along with precisely mapping TSSs in rat heart, we also show that a large number of genes use alternate promoters between the SHR and BN strains, suggesting a role of alternative promoter usage in complex disease.

## Results

### Identification of CAGE defined promoters in rat heart

To understand the role of alternate promoter usage and promoter shift in complex diseases, we performed CAGE tag sequencing from the left ventricles of the rat model of hypertension SHR and a normotensive rat strain BN in three replicates each from both male and female rats. We performed “non-amplifying, non-tagging Illumina CAGE” (nAnT-iCAGE) because no tagging protocol allows the sequencing of longer reads (100bp) and a non-amplification protocol eliminates biases due to PCR amplification^19^. On average we sequenced 15 million uniquely mapped read pairs per sample. To avoid mapping bias due to genomic variants between SHR and BN rat strains, we created a pseudo-SHR genome by substituting all the SHR single nucleotide variants (SNVs) in the BN reference genome^20^. CAGE tags from SHR were mapped to pseudo-SHR genome while BN CAGE tags were mapped to the BN reference genome. To obtain a comprehensive list of the rat heart promoters, we performed an analysis of all 12 samples together. We identified a total of 26,560 tag clusters, each representing a unique promoter region within the rat heart.

CAGE-defined promoters were annotated by overlapping them with the rat gene annotations from ENSEMBL (Version 93) ^21^. We defined a gene region as a region between the start and end positions of the longest transcript of a protein coding gene plus 1kb upstream region. Approximately, 80% (n=21,353) of the CAGE defined promoters were located in the gene regions of 10,935 protein coding genes (Figure 1A). We found that, in rat heart, a total of 6,445 genes use a single promoter, while 4,490 genes use more than one promoter (Supplemental Figure 1). For genes that use more than one promoter, the promoters were ranked based on their expression levels (TPM) as promoter 1 (P1), promoter 2 (P2) and so on.

**Figure 1:**
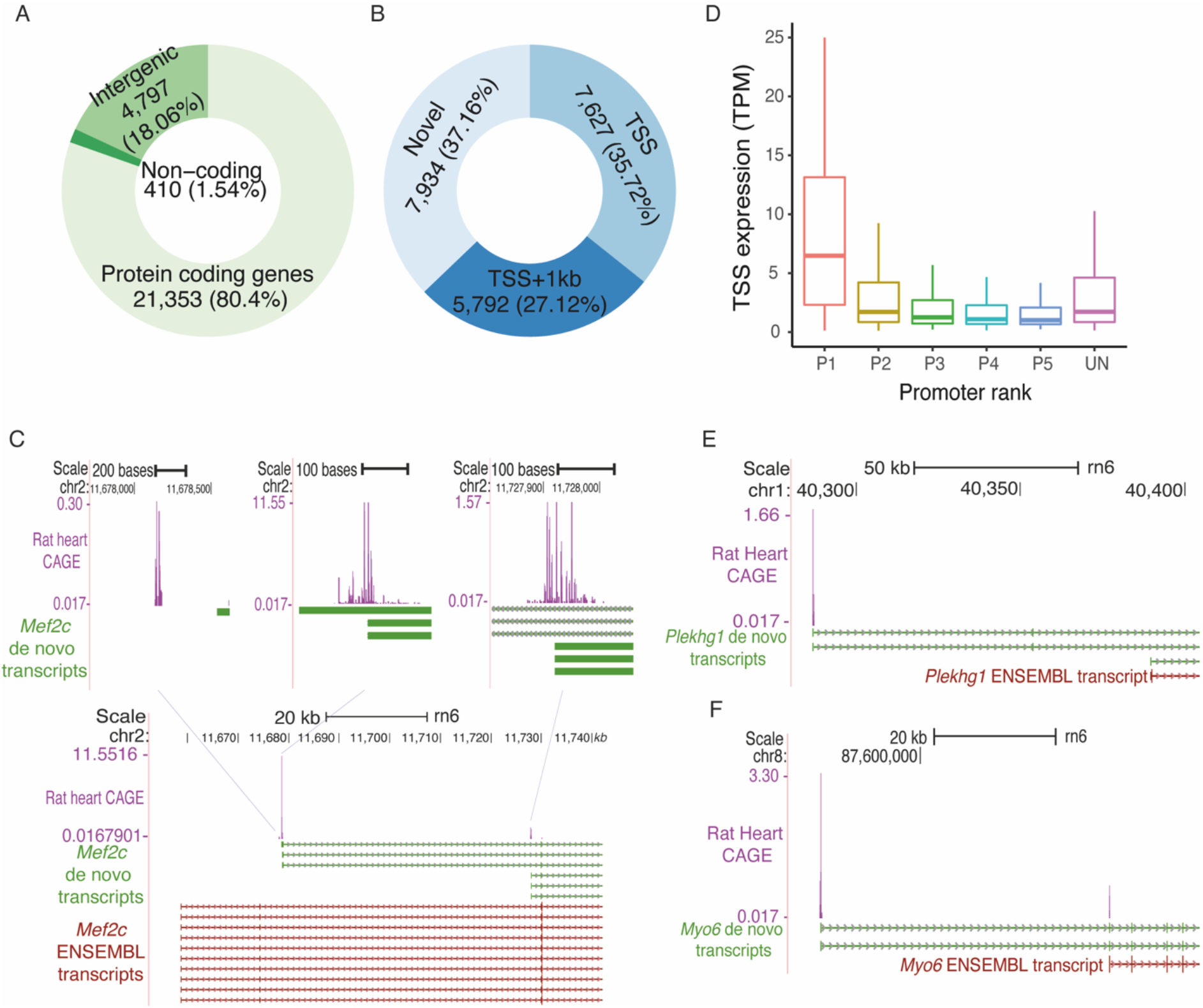
CAGE tag sequencing identifies promoters from rat left ventricle. **A.** Annotations of the CAGE defined promoters with respect to genomic features. **B.** Classification of CAGE defined TSS located in the gene regions, defined as the region between transcription start and end of the longest transcript of the gene plus 1kb upstream region, on same strand as that of the gene. **C.** Example of novel heart specific transcripts for identified using CAGE tag sequencing for gene *Mef2c*. **D.** Distribution of expression levels at promoters by promoter rank including the promoters that are unassigned to any protein coding gene. **E and F**. Examples of novel TSS annotations for gene *Plekhg1* and *Myo6* respectively.

### CAGE tag sequencing identifies novel cardiac TSS/transcripts

We evaluated CAGE defined TSS against the ENSEMBL TSS annotations for the rat genome (ENSEMBLE annotation version 93). Of 21,353 CAGE defined TSS that were within the gene regions, only 35.71% (n=7,627) matched perfectly with the ENSEMBL TSS annotations. Additional 1,965 tag clusters were within the 100bp of ENSEMBL annotated TSS, while 3,827 tag clusters were between 100bp to 1000bp away from the ENSEMBL annotated TSS. Surprisingly, no TSS annotation was found in ENSEMBL for the remaining 37% (n=7,934) tag clusters that were located within the protein coding gene regions and were on the same strand as that of the gene (Figure 1B). Most of these CAGE tag clusters might represent promoters of the novel cardiac transcripts. To investigate this hypothesis, we performed *de-novo* assembly of publicly available RNA-seq data from left ventricles of SHR and BN rats^16^. Of 7,934 tag clusters with no ENSEMBL TSS annotation, 383 overlapped perfectly with the TSS of *de-novo* transcripts, while 857 tag clusters were located within 500bp *de-novo* transcript TSS (Supplemental Table 1), suggesting that these tag clusters indeed represent promoters of novel heart specific transcripts.

A gene myocyte enhancer factor 2 (*Mef2c*), a member of MEF2 family of transcription factors, is a key regulator of cardiovascular development^22^. Loss of function (LoF) mutations in *MEF2C* have been implicated in dilated cardiomyopathy in human^23^. ENSEMBL annotations for *Mef2c* in rat genome show 10 different transcripts but all of them contain the same TSS (chr2:11,658,568). Interestingly, no CAGE tag cluster from rat heart overlap with the ENSEMBL annotated TSS of *Mef2c*. On the contrary, three CAGE defined tag clusters were located within the *Mef2c* gene body. The *de-novo* transcripts assembly identified six transcripts for *Mef2c* with three distinct TSSs, all of them overlapped with the CAGE defined TSSs (Figure 1C). The TSS of the *Mef2c* transcript that shows the highest expression (58.44 TPM) at the promoter was located 20kb downstream to the ENSEMBL annotated TSS. This suggest that our CAGE data could identify novel heart specific TSS of important cardiac genes.

A total of 5,207 tag clusters were located in intergenic regions of the genome, of which 410 were associated with annotated non-coding RNAs. We compared expression levels of the remaining 4,797 intergenic tag clusters with the expression levels of coding gene promoters ranked based on expression. The average expression levels of intergenic tag clusters ranged between the average expression of strongest promoter of genes (P1) and the second strongest promoter of the gene (P2; Figure 1D), suggesting that some of the intergenic tag clusters might also be promoters of the protein coding genes. To further investigate intergenic tag clusters, we compared them with the *de-novo* transcripts identified using RNA-seq data from rat heart. A total of 827 intergenic tag clusters were located within 100bp distance of *de-novo* transcript TSS. Additionally, 286 were within 500bp of *de-novo* transcript (Supplemental Table 2). For gene *Plekhg1*, no CAGE tag cluster was found at the ENSEMBL annotated TSS. The nearest CAGE tag cluster to gene *Plekhg1* was more than 100 kb upstream to ENSEMBL TSS, which was supported by our *de-novo* transcript assembly (Figure 1E). For the *Myo6* gene, only one transcript was annotated in ENSEMBL, TSS of which was supported by CAGE tag cluster. However, CAGE data and *de-novo* transcript assembly identified another transcript for the *Myo6* gene with a TSS around 47kb upstream of the ENSEMBL annotated TSS (Figure 1F). This novel transcript showed significantly higher expression (20.57 TPM) as compared to the ENSEMBL annotated transcript (0.96 TPM). Furthermore, the annotated transcript was upregulated in SHR as compared to BN while novel transcript was downregulated in SHR as compared to BN, suggesting alternative use of two *Myo6* transcripts in SHR and BN.

Here, we show that rat heart transcript annotations could be significantly improved using CAGE data by precisely mapping TSS and by identification of novel heart specific transcripts.

### CAGE identifies alternative promoter usage between SHR and BN

To understand the role of alternative promoter usage in disease we explored CAGE tag data from SHR and BN rat strains. We hypothesised that if a gene is predominantly transcribed from two different promoters in SHR and BN, respectively, then the tag clusters associated with both promoters should show differential expression but with expression difference in opposite direction. A total of 4,490 (41%) of the heart expressed genes showed more than one CAGE-defined promoter. We selected the promoters that showed a minimum of 20% expression of the total expression of all promoters of a gene. The majority (3,571) of the genes had only one predominant promoter, while 918 genes had at least two promoters with more than 20% activity. Of these 918 genes, for 471 genes none of the promoters show any difference in expression between SHR and BN, while for 447 genes at least one promoter showed statistically significant differential expression. For 420 genes all the promoters associated with a gene showed expression difference in same direction, while for 27 genes, at least two promoters showed differential expression in the opposite direction (Figure 2A). Suggesting that these 27 genes use alternative promoters between SHR and BN. For all these genes, use of alternative promoter between SHR and BN leads to a shorter transcript in one of the two strains with an impact on the protein coding region of the gene.

**Figure 2:**
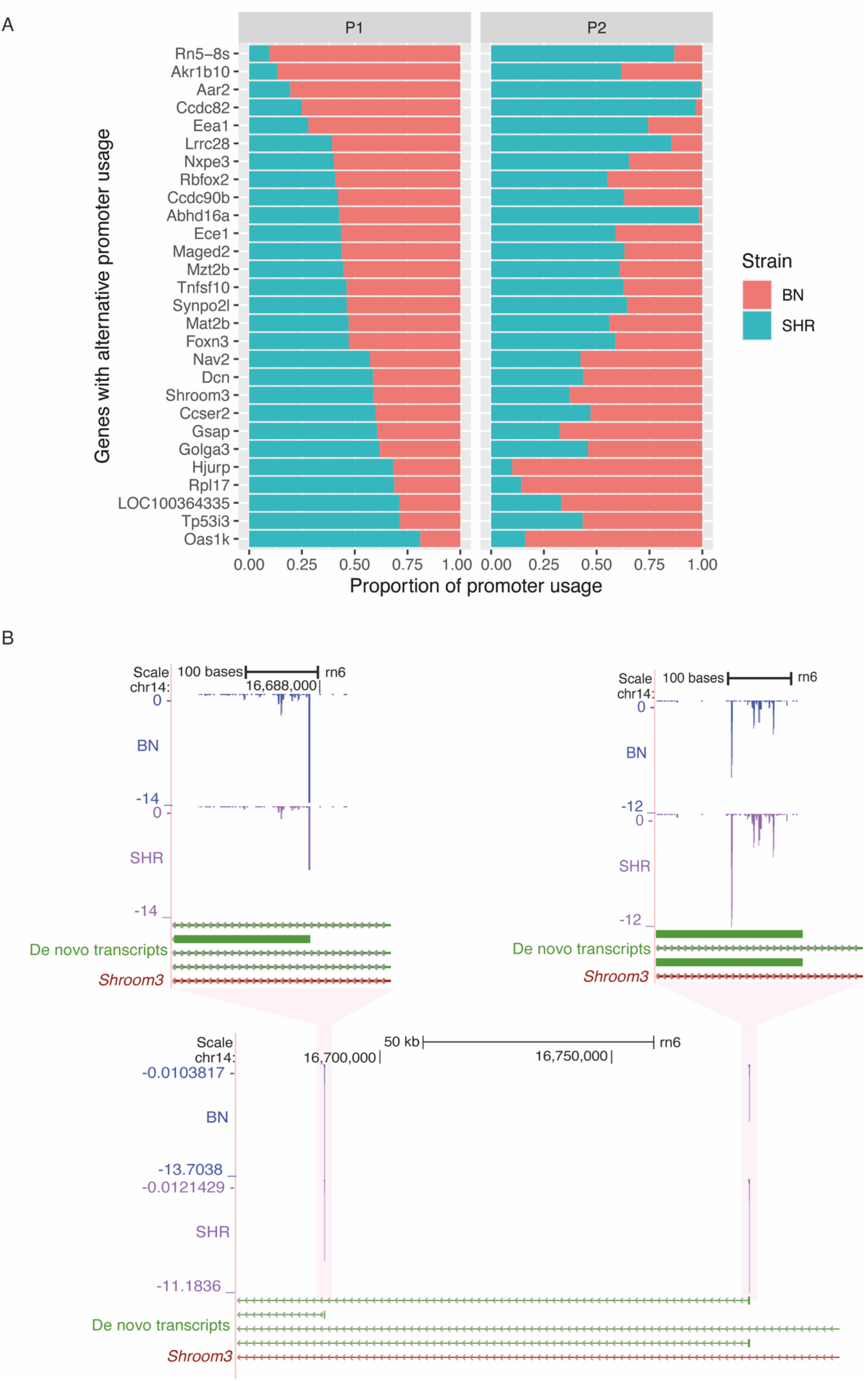
Alternative promoter usage between SHR and BN. **A.** list of genes with alternative promoter usage between SHR and BN. The P1 panel represents first promoter, while P2 panel represents second promoter. The x axis represents the proportion of the promoter used in SHR (green) and the BN (orange). **B**. Example of alternative promoter usage between SHR and BN in *Shroom3* gene. In SHR *Shroom3* is predominantly transcribed from the first promoter while in BN it is predominantly transcribed from the second promoter. The gene *Shroom3* is located on negative strand hence promoter expression levels have negative values.

These 27 genes include *Synpo2l*, *Abhd16a*, *Ece1*, *Shroom3* (Figure 2B) and *Rbfox2* all of which are implicated in cardiovascular disorders including hypertension^24–28^. Loss of function (LoF) mutations in *SHROOM3* and *SYNOP2L* genes have been shown to be associated with congenital heart defects^27^ and arterial fibrillation^24^ respectively. Dysregulation of *RBFOX2* has been shown to be an early event in cardiac pathogenesis of diabetes^28^. The polymorphisms in *ECE1* gene have been implicated in Human essential hypertension^26,29^, while *ABHD16A* is shown to be associated with coronary artery aneurism^25,30^.

### Promoter shift between SHR and BN

It has been shown that the switch in TSS usage within same promoter, often within 100bp, happens between maternal and the zygotic transcript during Zebrafish development^9^. We investigated whether TSS switching happens in a disease condition by using SHR and BN CAGE tag data. We identified 475 transcripts with a shift in TSS usage within the same promoter between SHR and BN, of which 287 (60%) were dominant promoters (P1) of the genes. However, only 50% of them were assigned to the ENSEMBL annotated genes while the remaining 50% were novel transcripts. In most cases, the TSS shift happened within 100bp of the same promoter, as observed previously^9^, and the shift happened in both directions with respect to reference rat strain (BN; Figure 3A). The ENSEMBL annotated genes that show shift in TSS usage between SHR and BN were enriched for genes involved in metabolic processes (*P*=0.003) and chloride transport (*P*=0.005). The genes that showed a switch in TSS usage within the same promoter between SHR and BN include, *Insr*, *Endog*, *Vnn1* and *Serpina3c*.

**Figure 3:**
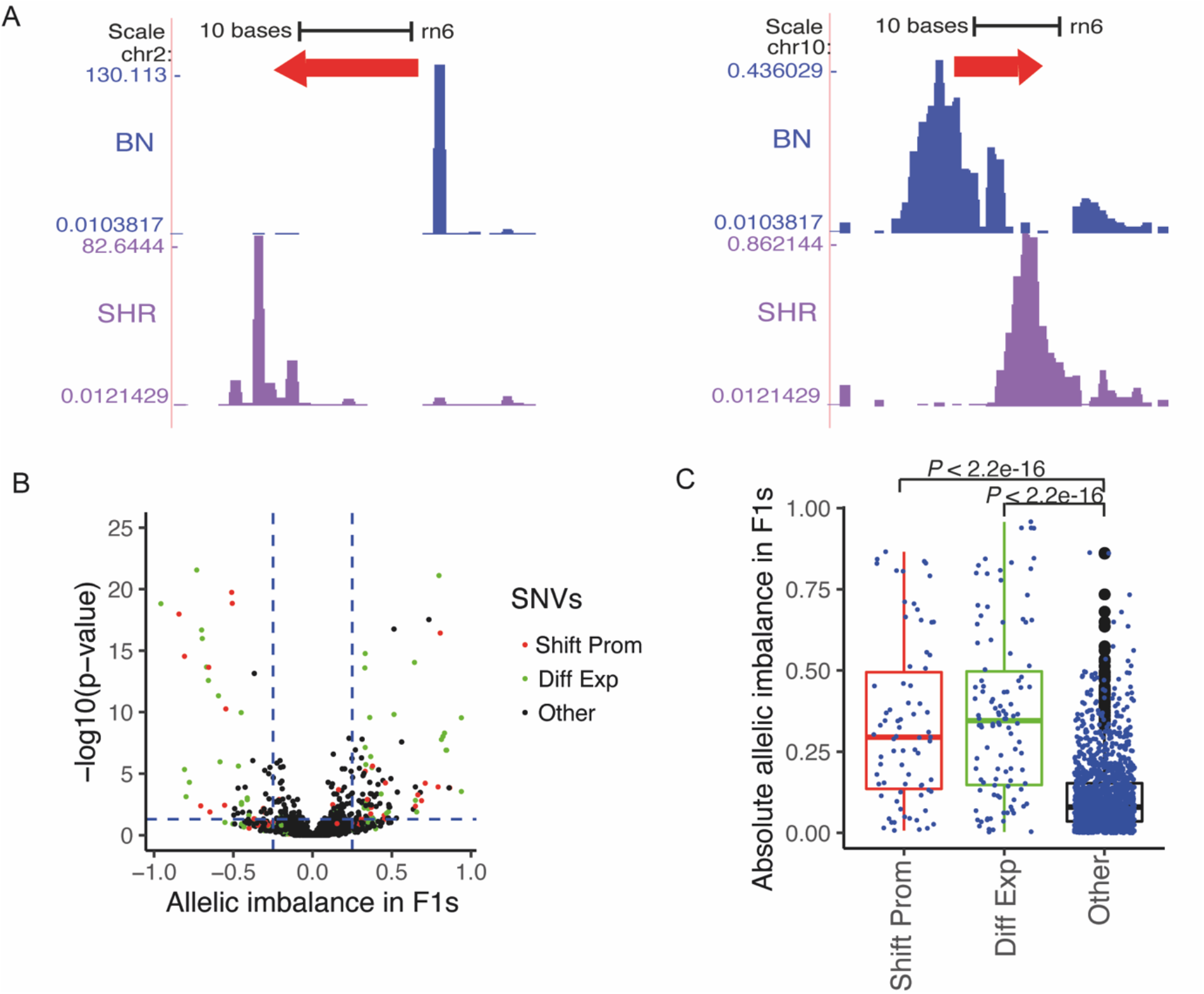
Shifting promoters between SHR and BN. **A**. Shift in TSS usage within same promoter happens in both directions with respect to reference BN strain. **B**. Allelic imbalance in F1 cross obtained by crossing SHR and BN. X axis represents allelic imbalance. The centre (zero) indicates no allelic imbalance between SHR and BN allele in F1. Positive values indicate allelic imbalance towards in reference (BN) allele while negative values indicate allelic imbalance towards SHR allele. Horizontal blue dotted line corresponds to p-value equal to 0.05 while vertical blue dotted line corresponds to allelic imbalance of 0.25 and -0.25. The red dots represent variants located in shifting promoters while green dots represent variant in promoters that show differential expression between parental strains (SHR and BN). Black dots represent genomic variants in CAGE defined promoters that are neither shifting promoters nor differentially expressed between parental strains. **C**. Box plot showing distribution of allelic imbalance in F1 cross in shifting promoters, differentially expressed promoters and remaining CAGE defined promoter regions. Shifting promoters (red box) show significantly (Mann-Whitney test) higher proportion variants with allelic imbalance as compared to non-shifting and non-differentially expressed promoters (black box). Shift Prom: Shifting promoters, Diff Exp: Differentially expressed promoters in parental strains (SHR and BN), Other: CAGE defined promoters that are neither shifting promoters nor differentially expressed in parental strains.

The transcripts in which the shift in TSS usage between two rat strains was observed, produce a longer transcript in one strain comparted to a shorter transcript in another, in most cases affecting the length of the 5’ untranslated region (UTR). Being inbred rat strains, SHR and BN are completely homozygous, hence the F1 cross between SHR and BN rat strains tend to be heterozygous at all the variable loci between SHR and BN. Thus, F1s will inherit both longer and shorter transcripts of the genes that show promoter switching between the two rat strains. We reasoned that the genomic variants located in regions that are unique to longer transcripts should show strong allelic imbalance in F1s. To test this hypothesis, we generated reciprocal F1 crosses by crossing SHR females with BN male and BN female with SHR male. For each reciprocal cross, we performed CAGE tag sequencing of LVs from three male and three female rats, totalling 12 F1 rats.

A total of 1,853 genomic variants between SHR and BN were located in CAGE tag clusters (CAGE defined promoters), of which 110 variants were located in shifting promoters, remaining in normal promoters. The variants located in shifting promoters showed strong allelic imbalance in F1s (Figure 3B) as compared to variants located in the non-shifting, non-differentially expressed promoters (Fisher’s exact test, *P* < 2.2e^-16^). In F1s, allelic imbalance in shifting promoters was comparable to allelic imbalance of variants located in promoters that were strongly differentially expressed (adjusted p-value< 0.05 and log2 fold change > 1) in parental strains (SHR and BN; Figure 3C). As expected, the variants located in non-differentially expressed promoters did not show allelic imbalance (Figure 3B,C). The strong allelic imbalance of variants located in shifting promoters confirms the presence of shifting promoters between SHR and BN rat strains.

### Promoter shift in *Insr* gene

The insulin signalling pathway plays a key role in metabolic regulation, growth control and neuronal function^31^. The insulin signalling pathway is mediated by the insulin receptor (*Insr*), a transmembrane receptor with tyrosine kinase activity^32^. We found that *Insr* shows a shift in TSS usage between SHR and BN (Figure 4A). In the BN rat strain, transcription started predominantly from genomic position chr12:1,816,432, which was 346bp upstream to the *Insr* start codon (chr12:1,816,086), while in SHR, the TSS was observed 414bp upstream (chr12:1,816,500) to the *Insr* start codon, resulting in longer transcript in SHR as compared to BN. To confirm this finding, we investigated RNA-seq data from SHR and BN heart. No RNA-seq reads were observed in BN heart in the extended region that was specific to SHR, supporting the findings from CAGE data (Figure 4A).

**Figure 4:**
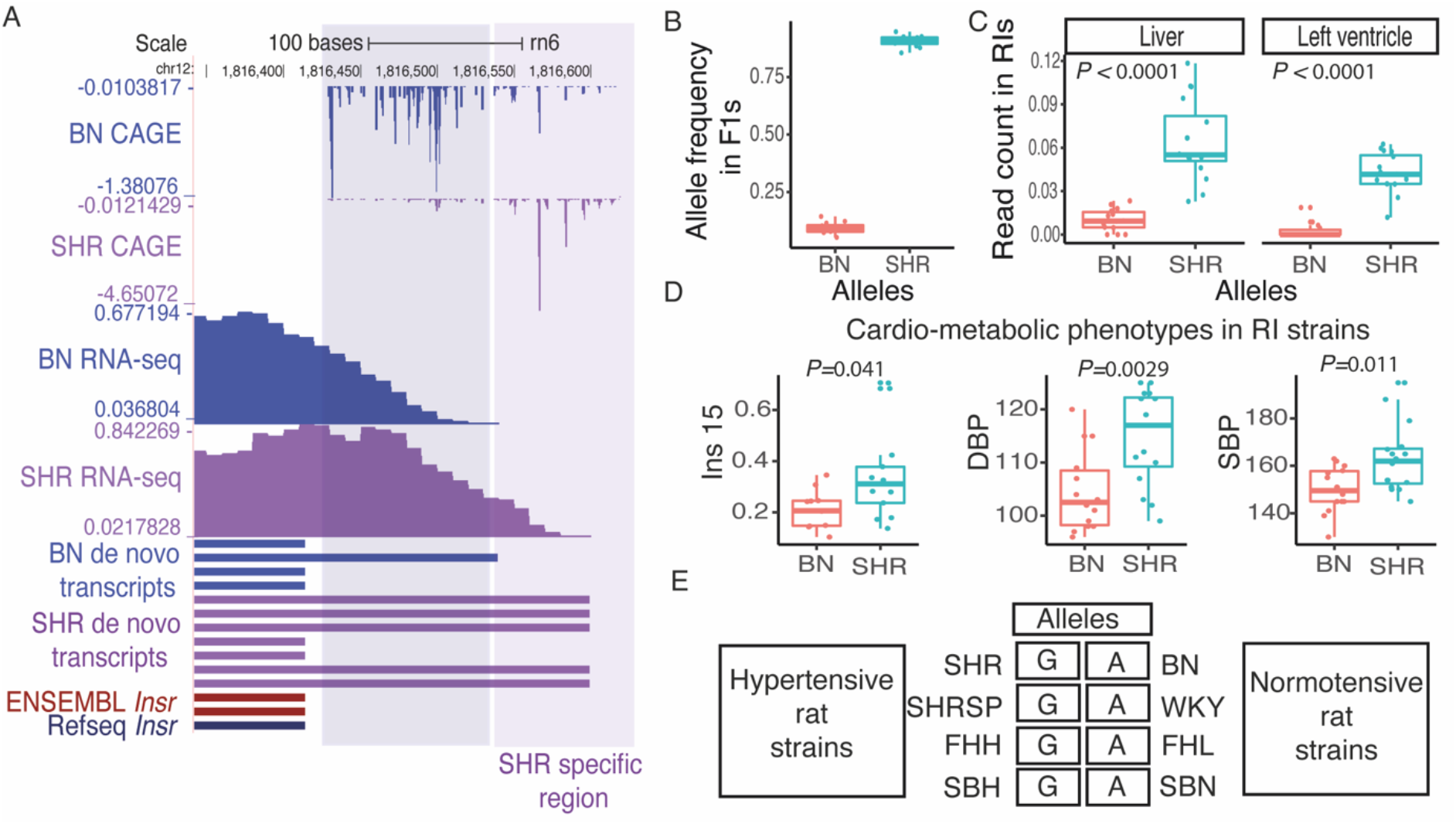
Shifting promoter in insulin receptor (*Insr*) gene. **A.** Shifting promoter in *Insr*. The GAGE tag sequencing identified promoter shift between SHR and BN strain for *Insr* gene which leads to longer *Insr* transcript in SHR as compared to BN. Finding from CAGE was confirmed by RNA-seq data from rat left ventricle as well as *de novo* transcript assembly generated using rat left ventricle RNA-seq data. **B**. The variant g.1816552:A>G located in SHR specific *Insr* transcript region shows significant allelic imbalance in F1 cross with majority of the reads showing SHR allele in F1s. **C.** Based on the genotype at g.1816552:A>G recombinant inbred (RI) strains were grouped into two groups, where AA represents RI strains inheriting BN allele while GG represents RI strains inheriting SHR allele. RNA-seq read count in SHR specific *Insr* transcript region were normalised to the read count from first exon of *Insr* gene using publicly available RNA-seq data from left-ventricle and liver in RI strains. Both in left ventricle and liver, RI strains that inherited SHR allele showed higher normalised reads count suggesting that they harbour longer transcript for *Insr* gene, while RI strains that inherited BN allele use shorter transcript. **D.** RI strains that harbour SHR allele showed significantly higher levels of insulin, systolic and diastolic blood pressure. **E**. Genotype in rat models of hypertension and their control strains at variant (g.1816552:A>G) located in SHR specific *Insr* transcript region. All hypertensive rat strain contains GG allele, same as SHR while all the normotensive rat strains contain AA allele, same as BN.

A genomic variant (g.1816552:A>G) was located in the SHR specific region of *Insr* transcript. The variant (g.1816552:A>G) showed significant allelic imbalance (Binomial test, *P*=2.77e^-15^) in F1 cross with the majority of reads overlapping the SHR specific region showed in the SHR allele at the variant position (Figure 4B) suggesting that the longer transcript was inherited from SHR in F1s. This finding confirms the shift in TSS usage between SHR and BN for the gene *Insr* in heart. The HXB/BXH panel of recombinant inbred (RI) rat strains generated by crossing SHR and BN have been widely used for genetic mapping^33^. Recently, RNA-seq has been performed in heart and liver tissues from RI strains^34^. Using the variant g.1816552:A>G we grouped RI strains into two groups, one that inherited SHR allele and the other BN allele. The RI strains that inherited the SHR allele predominantly used a longer transcript both in heart and Liver (Figure 4C).

Next, we investigate the cardio-metabolic phenotypes measured in RI strains^35,36^. We found significant association between the RI strain genotype at g.1816552:A>G and the insulin level^35^ (Mann-Whitney test*, P*= 0.041), systolic blood pressure^36^ (Mann-Whitney test, *P*=0.0029) and diastolic blood pressure^36^ (Mann-Whitney test, *P*=0.011) in RI strains. The RI strains containing the SHR allele (GG; long transcript) showed significantly elevated levels of insulin, systolic and diastolic blood pressure (Figure 4D). Interestingly, a large number of physiological quantitative trait loci (pQTLs) for blood pressure and insulin resistance have been mapped in the region harbouring *Insr* (Rat Genome Database; http://rgd.mcw.edu/).

Multiple rat strains have been developed to study hypertension and the whole genome of most of the rat models of hypertension have been sequenced^17^. We found that multiple rat models of hypertension; Fawn Hooded Hypertensive (FHH), Sabra Hypertensive (SBH) and Spontaneously Hypertensive Rat Stroke Prone (SHRSP), also harbour SHR allele at *Insr* promoter while respective control strains Fawn Hooded Low (FHL) blood pressure, Sabra Normotensive (SBN) and Wistar Kyoto (WKY) contain BN allele (Figure 4E). Suggesting strong association between longer *Insr* transcript and the cardio-metabolic phenotype shown by these rat strains.

Taken together, multiple lines of genetic evidence suggests that the switch in TSS usage for insulin receptor (*Insr*) gene in SHR might be associated with the disease phenotypes observed in SHR rat strain. However, strong experimental support is required to establish causal relationship between differential *Insr* TSS usage in SHR and the disease phenotype.

## Discussion

Cap analysis of gene expression (CAGE) has been widely used to precisely map transcription start sites (TSS) in humans and mouse^4,37^. Using CAGE it has been shown that the promoter usage tends to be tissue and developmental stage specific^4,9^. Ubiquitously expressed genes achieve cell type specificity through use of cell type specific transcription start sites^38^. Though the role of alternate promoter usage has been well established in tissue specific expression of genes, its role in disease has been poorly understood. To understand the role of alternate promoter usage in complex disease we performed CAGE tag sequencing of left ventricles of two rat strains, SHR a widely used model to study hypertension and normotensive rat strain BN, progenitors of BXH/HXB RI strains.

Rat is a widely used animal model to study various disease phenotypes specifically cardio-metabolic phenotypes^12^, however, to date the precise TSS for rat genes have not been mapped. In this study, we show that CAGE tag sequencing significantly improves the TSS annotations of rats. Using combined analysis of *de-novo* transcript assembly and CAGE tag sequencing we show that for 1,113 transcripts TSS was located more than 1000 bp upstream to the annotated TSS. In addition, we identified a large number of cardiac TSSs that were not annotated in ENSEMBL, this included three novel TSSs for myocyte enhancer factor 2 (*Mef2c*). In this study, we could identify thousands of novel TSS/transcripts even though we performed CAGE tag sequencing from only one rat tissue. This suggests that CAGE tag sequencing from wide range of rat tissues could significantly improve rat TSS/transcript annotations. In addition, it will help to understand tissue specific usage of TSS in rat and it would facilitate comparative analysis with human and mouse CAGE data.

To understand the role of alternative promoter usage in disease we used CAGE tag data from SHR and BN rat strains. We identified two types of TSS switching events between SHR and BN. First, where gene use completely different promoters between two rat strains that are locate more than 100bp away from each other and affect the coding region of the gene. This even leads to shorter protein sequence due to truncated N terminus in one strain compared to other. Both the transcripts show expression in both strains; however, one transcript tends to be predominantly expressed in one strain while other transcript show predominant expression in the other strain. We identified 27 genes with alternative promoter usage between SHR and BN, a proportion of them were known to be associated with cardiovascular disorders. This suggests that alternative promoter usage could potentially result in complex disease phenotype. Experimental validation is required to confirm these findings; however, this study could be the first step towards understanding role of alternative promoter usage in complex disorders such as hypertension.

In the second type of TSS switching event, transcripts use different TSS within same promoter. In most cases the TSS switch happened within 100bp of each other. The TSS switching event does not affect protein coding regions of the gene but they alter the length of 5’ untranslated regions (UTR). 5’ UTR plays a major role in translational efficiency as 5’UTR is critical for ribosome recruitment to mRNA choice of start codon^39^. The 5’ UTR contain key elements of translational regulation including structural motifs and uORFs^40^. These sequence elements in the 5’ UTR control contribute to mRNA translation by controlling selection of translational initiation sites (TIS)^39,40^. Sequence variation even in only 10bp immediate upstream of translational start sites or uORFs lead to profound changes in translational efficiency and in the amount of protein produced^40,41^. Thus, a switch of TSS usage between two strains even within 100bp could have profound impact on the protein translation from mRNA. In this study, we identified 425 transcripts with TSS switching between SHR and BN. Furthermore, we show that the genomic variants located in these regions show significant allelic imbalance in F1s derived by reciprocal cross of SHR and BN, confirming TSS switch. TSS switching between SHR and BN might lead to complex disease phenotypes shown by SHR, however, further experimental studies are required to establish causal role of TSS switching in disease manifestation.

We identified TSS switching in insulin receptor (*Insr*) gene between SHR and BN rat strain resulting in longer transcript with extended 5’ UTR in SHR compared to BN. The genomic variant located in SHR specific region showed significant allelic imbalance in F1s, confirming the TSS switch. Furthermore, *Insr* TSS usage was strongly associated with the insulin levels, systolic and diastolic blood pressure in HXB/BXH panel of recombinant inbred (RI) strains derived by crossing SHR and BN rat strains. The SHR rat strain shows various phenotypes such as hypertension, insulin resistance, dyslipidaemia, and central obesity, collectively known as metabolic syndrome^42^. The components of metabolic syndrome, hypertension, and insulin resistance often coexist^43,44^. Clinical studies have shown that 50% of the hypertensive individuals have hyperinsulinemia while 80% of type 2 diabetes patients have hypertension^44^. Previous studies suggest the causal relationship between compensatory hyperinsulinemia due to insulin resistance and hypertension^44,45^. It is challenging to establish causal relationship between *Insr* TSS switch and disease phenotypes shown by SHR, however, multiple lines of genetic evidence along with the strong association between *Insr* TSS usage with insulin levels and blood pressure in RI strains suggest that alternate TSS usage might have phenotypic impact in SHR rat strain.

Taken together our study suggests that alternative promoter usage may play a role in complex disease phenotypes and paves the way for further experimental studies to establish a causal relationship between alternative promoter usage and complex diseases. Furthermore, data generated in this study could help in improving rat gene annotations, which would significantly benefit researchers who use rat as a model organism to study complex diseases.

## Materials and Methods

### Rat strains

The rat model of hypertension, Spontaneously Hypertensive Rat (SHR/OlaIpcv) and the Brown Norway (BN-Lx/Cub) rat strains (referred as a SHR and BN) were maintained in animal facility at the Institute of Physiology, Czech Academy of Sciences, Prague, Czech Republic. All the experimental procedures were carried out as per European Union National Guidelines and the animal protection laws of the Czech Republic and were approved by the ethics committee of the Institute of Physiology, Czech Academy of Sciences, Prague.

Left ventricles from six week old SHR (n=6, three males and 3 females) and BN (n=6, three males and 3 females) were harvested at the Institute of Physiology, Czech Academy of Sciences, Prague. Reciprocal F1 cross were generated by crossing SHR females with BN males (SHRxBN) and BN females with SHR males (BNxSHR). Left ventricles from six week old F1s were harvested. We used total 12 F1s with six (three males and three females) F1s from each reciprocal cross.

### CAGE tag sequencing

The left ventricles harvested from rat strains were sent to DNAform, Japan. The RNA extraction and non-tagging, non-amplification (nAnT)-CAGE tag sequencing was performed at DNAform, Japan following a previously described protocol^19^.

### Read mapping

The CAGE tags were sequenced using 100bp paired end sequencing technology on Illumina HiSeq2500 platform. The adaptors were removed using cutadapt^46^. The additional G nucleotide, which is often attached to the 5’ end of the tag by the templet free activity of the reverse transcriptase in the cDNA preparation step of the CAGE protocol, was removed. The CAGE tag sequences were mapped to reference genome using STAR-2.4.0^47^. To avoid read mapping bias due to genomic variants between the SHR and BN rat strains, a pseudo-SHR genome was generated by substituting all SHR SNVs^10,17^ from BN reference genome. CAGE tags from SHR LVs were mapped to the pseudo-SHR reference genome, while BN LV CAGE tags were mapped to the BN reference genome^20^. CAGE tags from F1s were mapped to both BN and pseudo-SHR reference genomes and the best hits were selected for the downstream analysis.

### Identification of CAGE defined tag clusters

All the replicates from parental strains (SHR and BN) were used to identify CAGE defined tag clusters (TC). Only uniquely mapped reads, reads mapped with quality score 255 extracted using samtools-1.2^48^, were used to identify CAGE defined tag clusters. All the unique 5’ ends of first read (R1) of the read pair were considered as a CAGE defined TSS (CTSS) and counts were generated for each position representing a number of unique tags starting from that position. All the positions where at least three samples showed minimum one read count were selected. The CTSSs that were within 20bp distance were merged to generate TC. Counts within each tag cluster were then normalised to one million reads (TPM). The TCs that had normalised read count of 1TPM in at least one sample were selected as a final set. Next, the TCs that were within 100bp away from each other were clustered together to define the promoter regions.

### Promoter Ranking

The gene regions were defined as a genomic region between start and end position of the longest transcript of a gene plus 1000bp upstream of the start position. All the CAGE-defined promoters located within the gene regions on the same strand were assigned to the gene. The promoters were then ranked based on the expression levels (TPM) in descending order. The promoter that showed the highest expression was called P1 and subsequent promoters were called P2, P3 and so on for each gene. All the tag clusters that were located in intergenic regions of the genome were considered as independent promoters with P1 rank.

### Identification of genes with alternate promoter usage

To identify genes with alternate promoter usage between SHR and BN, first differentially expressed tag clusters were identified using DEseq2-1.32.0^49^. The promoters that were differentially expressed (adjusted p-value <=0.05) between SHR and BN were extracted. All the promoters that showed at least 20% expression compared to the aggregate expression levels of the all the promoters of the same gene were selected for the downstream analysis. The genes where two or more promoters showed differential expression in opposite direction, i.e. of the two promoters of the same gene, one promoter showing upregulation in SHR compared to BN while another showing downregulation in SHR as compared to BN, were selected. A gene was called to use alternate promoters only when two promoters that showed expression difference in opposite direction included P1 promoter.

### Identification of shifting promoters

To identify switching promoters, for each TC (promoter) the CTSS data (count of unique 5’ ends of CAGE tags within TC at base pair resolution) from six replicates of each parental strains (SHR and BN) was aggregated. The Kolmogorov–Smirnov test was used to identify switching promoters. Following filtering criterias were used to select statistically significant promoter switching events between SHR and BN. i) The test statistics *D* must be greater than 0.3. ii) The test statistics *D* must be greater than the critical value at alpha 0.05. The genomic position where the highest *D* score was achieved was selected as the position of switch.

### RNA-seq data analysis

The publicly available RNA-seq data from SHR and BN left ventricle and liver^16^ was downloaded from SRA/ENA. Adaptors were removed using cutadapt. RNA-seq reads from the BN were mapped to the BN reference genome while SHR RNA-seq reads were mapped to the pseudo-SHR genome using STAR-2.4.0^47^.

### *De novo* transcriptome assembly

*De novo* transcriptome assembly was performed using stringtie-2.1.4^50^ with default parameters using mapped RNA-seq read files (bam files) generated by STAR-2.4.0^47^. *De novo* assembly was performed independently for SHR and BN transcriptome. In addition, to generate *de novo* transcripts for rat heart, transcriptome assemblies from SHR and BN heart were merged using stringtie-2.1.4 merge.

### Allelic imbalance analysis in F1s

Variants in CAGE derived tag clusters were identified using GATK-4.0.7^51,52^ following the recommended guidelines for variant calling from transcriptome data. Reads that spanned intronic region are mapped to exons by splitting reads. To avoid calling variants in such regions, split reads spanning intronic regions were pre-processed using GATK SplitNCigarReads. Then variants were identified in each sample independently using GATK HaplotypeCaller. The gVCF files were then merged using GATK CombineGVCFs.

Following filtering criteria were used to select variants for allelic imbalance analysis. i) variants must be covered with minimum 10 reads in all six replicates of both the parental strain. ii) variants must be variable between the two parental strains SHR and BN. iii) In F1, variants must be covered with minimum 10 reads in at least nine out of 12 replicates.

To identify the variants that show allelic imbalance in F1s, reference (BN) allele and alternate (SHR) allele counts were extracted for each high-quality variant. A binomial test was used to identity variants with allelic imbalance in F1s. The variants that showed p-value <= 0.05 were called as allelically imbalanced variants in F1s.

### RNA-seq data from RI strains

The left ventricle and liver RNA-seq data from RI strains was obtained from previous publication^53^.

### *Insr* promoter analysis in RI strains

Based on the genotype at g.1816552:A>G recombinant inbred (RI) strains were grouped into two groups, where AA represents RI strains that inherited the BN allele while GG represents RI strains inherited the SHR allele. RNA-seq read count were obtained from the SHR specific *Insr* transcript region (chr12:1,816,550-1,816,650). The RNA-seq read count from the SHR specific *Insr* transcript region was then normalised to the read counts from first exon of the *Insr* gene (chr12:1,815,967-1,816,414). The Mann-Whitney test was used to identify significant association between genotype and normalised read count in SHR specific *Insr* promoter region.

### Phenotypic measurements in RI strains

The phenotypic measurements such as insulin levels, systolic and diastolic blood pressure in RI strains were obtained from previous publications^35,36^. The Mann-Whitney test was used to identify significant association between genotype and the phenotype in RI strains.

### Gene enrichment analysis

Gene enrichment analysis was performed using Enrichr web browser (https://maayanlab.cloud/Enrichr/).

